# The enrichment of self-domestication and neural crest function loci in the heritability of neurodevelopmental disorders is not independent of genomic regulatory functions

**DOI:** 10.1101/2022.09.26.509526

**Authors:** Dora Koller, Antonio Benítez-Burraco, Renato Polimanti

## Abstract

Self-domestication could play an important role in contributing to shape the biology of human brain and the predisposition to neurodevelopmental disorders. Leveraging genome-wide data from the Psychiatric Genomics Consortium, we tested the enrichment of self-domestication and neural crest function loci with respect to the heritability of autism spectrum disorder, schizophrenia (SCZ in East Asian and European ancestries, EAS and EUR, respectively), attention-deficit/hyperactivity disorder, obsessive-compulsive disorder, and Tourette’s syndrome (TS). Considering only self-domestication and neural-crest-function annotations in the linkage disequilibrium score regression (LDSC) model, our partitioned heritability analysis revealed statistically significant enrichments across all disorders investigated. The estimates of the heritability enrichments for self-domestication loci were similar across neurodevelopmental disorders, ranging from 0.902 (EAS SCZ, p=4.55×10^−20^) to 1.577 (TS, p=5.85×10^−5^). Conversely, a wider spectrum of heritability enrichment estimates was present for neural crest function with the highest enrichment observed for TS (enrichment=3.453, p=2.88×10^−3^) and the lowest for EAS SCZ (enrichment=1.971, p=3.8l×10^−3^). Although these estimates appear to be strong, the enrichments for self-domestication and neural crest function were null once we included additional annotations related to different genomic features. This indicates that the effect of self-domestication on the polygenic architecture of neurodevelopmental disorders is not independent of other functions of human genome.

---

Self-domestication could play an important role in contributing to the human phenotypic spectrum. According to this hypothesis, our species experienced an evolutionary process similar to animal domestication that paved the way toward enhanced social cognition, increased cooperation, and extended social networks, which led to our advanced technology and sophisticated culture(Hare 2017; Thomas and Kirby 2018). Also, the emergence of modern languages has been implicated with the self-domestication hypothesis, where the reduction in reactive aggression – a key factor in self-domestication processes – enabled humans to exploit cognitive abilities and social behaviors to evolve more sophisticated grammars, as well as a specific form of communication governed by persuasive reciprocity (Thomas and Kirby 2018; Benitez-Burraco and Progovac 2020; Benitez-Burraco, Ferretti, et al. 2021). In animals, selection for tameness, the core of all domestication events, results in a constellation of physical, cognitive, and behavioral traits (the domestication syndrome). This appears to be due to the effect of tameness selection on the input to the neural crest, an embryonic structure giving rise to many different body parts during development (Wilkins, et al. 2014; Sanchez-Villagra, et al. 2016; Lord, et al. 2020). A similar evolutionary process could have also happened in our species in response to external factors (e.g., climate deterioration during the last glaciation or changes in our ecology), promoting selection of less emotionally reactive and more prosocial individuals. In previous paleo-genomic studies, there were significant positive-selection signatures between anatomically modern humans and several domesticated species (Theofanopoulou, et al. 2017; Benitez-Burraco, Chekalin, et al. 2021). The overlapping genes under positive selection in both anatomically modern humans and domesticated species were enriched for signaling pathways and cell lineages, also including the neural crest (Theofanopoulou, et al. 2017). One intriguing question is whether self-domestication also contributes to non-neurotypical human phenotypes, so that abnormal self-domestication can be regarded as one etiological factor of psychiatric disorders and altered behavioral traits within the human species. Genome-wide association statistics generated by large-scale studies of psychiatric disorders and traits have been investigated with respect to the possible SNP-based heritability (SNP-h2) enrichments for loci affected by different evolutionary mechanisms such as positive and negative selection, and Neanderthal introgression. Although enrichment for positive selection and Neanderthal introgression signatures in the genetic liability to certain psychiatric disorders such as schizophrenia (SCZ) (Srinivasan, et al. 2016) and autism spectrum disorders (ASD) (Polimanti and Gelernter 2017), further studies showed that these enrichments were likely due to the unaccounted presence of negative selection signature and the effect of other regulatory functions (Pardinas, et al. 2018; Wendt, et al. 2021). Indeed, the application of the linkage disequilibrium score regression to perform a stratified heritability analysis permitted to estimate SNP-h2 enrichments by assessing multiple genomic annotations simultaneously (Finucane, et al. 2015). To our knowledge, a similar systematic analysis has not been conducted to test the self-domestication hypothesis in the context of the genetic liability to psychiatric disorders, despite some evidence supporting this possibility. Features of self-domestication have been found altered in people with ASD (Benitez-Burraco, et al. 2016) and SCZ (Benitez-Burraco, et al. 2017). Furthermore, loci implicated in mammal domestication appear to be dysregulated in the blood of individuals with ASD (Benitez-Burraco 2020), whereas genes exhibiting methylation changes in ASD and SZ are enriched in processes such as neural crest differentiation and ectoderm differentiation (Anastasiadi, et al. 2022). Likewise, a previous study showed the polygenic component of psychiatric disorders and behavioral traits is associated with the phonological complexity of European languages in relation to local adaption (Polimanti, et al. 2018), with types of languages being impacted, as noted, by changes in self-domestication levels (Benitez-Burraco and Progovac 2020). These findings suggest that evolutionary pressures implicated in human self-domestication processes may have contributed to shaping the polygenic architecture of ASD, SCZ, and other neurodevelopmental disorders. Exploring this aspect could generate new insights into human evolutionary history and the etiology of high prevalent cognitive conditions.

To assess systematically the possible contribution of loci implicated in self-domestication and neural crest function, we leveraged genome-wide association statistics generated from large-scale studies conducted by the Psychiatric Genomics Consortium (Table 1). We focused our attention on five psychiatric disorders, including ASD (Grove, et al. 2019), SCZ in East Asian and European ancestries (EAS and EUR, respectively; (Trubetskoy, et al. 2022), attention-deficit/hyperactivity disorder (ADHD) (Demontis, et al. 2019), obsessive-compulsive disorder (OCD) (International Obsessive Compulsive Disorder Foundation Genetics and Studies 2018), and Tourette’s syndrome (TS) (Yu, et al. 2019).

**Table 1.** Details of the genome-wide association studies investigated in the present study.

| Disorder | Abbreviation | $h^2$ (se)<sup>*</sup> | N<sub>cases</sub> | N<sub>controls</sub> | N<sub>eff</sub> |
| --- | --- | --- | --- | --- | --- |
| Attention-deficit/hyperactivity disorder (Demontis, et al. 2019) | ADHD | 0.22 (0.01) | 20,183 | 35,191 | 51,306 |
| Autism spectrum disorders (Grove, et al. 2019) | ASD | 0.12 (0.01) | 18,381 | 27,969 | 44,366 |
| Obsessive-compulsive disorder (International Obsessive Compulsive Disorder Foundation Genetics and Studies 2018) | OCD | 0.28 (0.04) | 2,688 | 7,037 | 7,780 |
| Schizophrenia (European ancestry) (Trubetskoy, et al. 2022) | SCZ (EUR) | 0.24 (0.02) | 53,386 | 77,258 | 126,281 |
| Schizophrenia (East Asian Ancestry) (Trubetskoy, et al. 2022) | SCZ (EAS) | 0.23 (0.03) | 14,004 | 16,757 | 30,514 |
| Tourette's syndrome (Yu, et al. 2019) | TS | 0.21 (0.02) | 4,819 | 9,488 | 12,783 |

Abbreviations: ADHD: attention deficit-hyperactivity disorder, ASD: autism spectrum disorder, OCD: obsessive compulsive disorder, SCZ: schizophrenia and TS: Tourette's syndrome, N<sub>cases</sub>: number of cases, N<sub>controls</sub>: number of controls, N<sub>eff</sub>: effective sample size. \*SNP-based heritability ( $h^2$ ) and standard error (se) on the liability scale.

Following the guidelines to perform a stratified SNP-h2 analysis using LDSC method (Finucane, et al. 2015), we created two binary annotations related to loci involved in self-domestication and neural crest function. The loci related to self-domestication were derived from genes positively selected in multiple domesticated mammals (Supplemental Table 1). The loci involved in neural crest development and function were identified considering multiple data sources, such as genes associated with neurocristopathy and genes involved in neural crest induction, specification, signaling, and differentiation (Supplemental Table 2). A detailed description of the approaches and the data sources is provided in the Material and Methods section.

Including only the annotations for self-domestication and neural crest function in the LDSC model, we observed enrichments surviving false discovery rate multiple testing correction (FDR q<0.05) for both across all the disorders investigated (Figure 1; Supplemental Tables 3 and 4) with the exception of OCD with respect to neural crest function (Enrichment=1.972, p=0.343).

**Figure 1.**
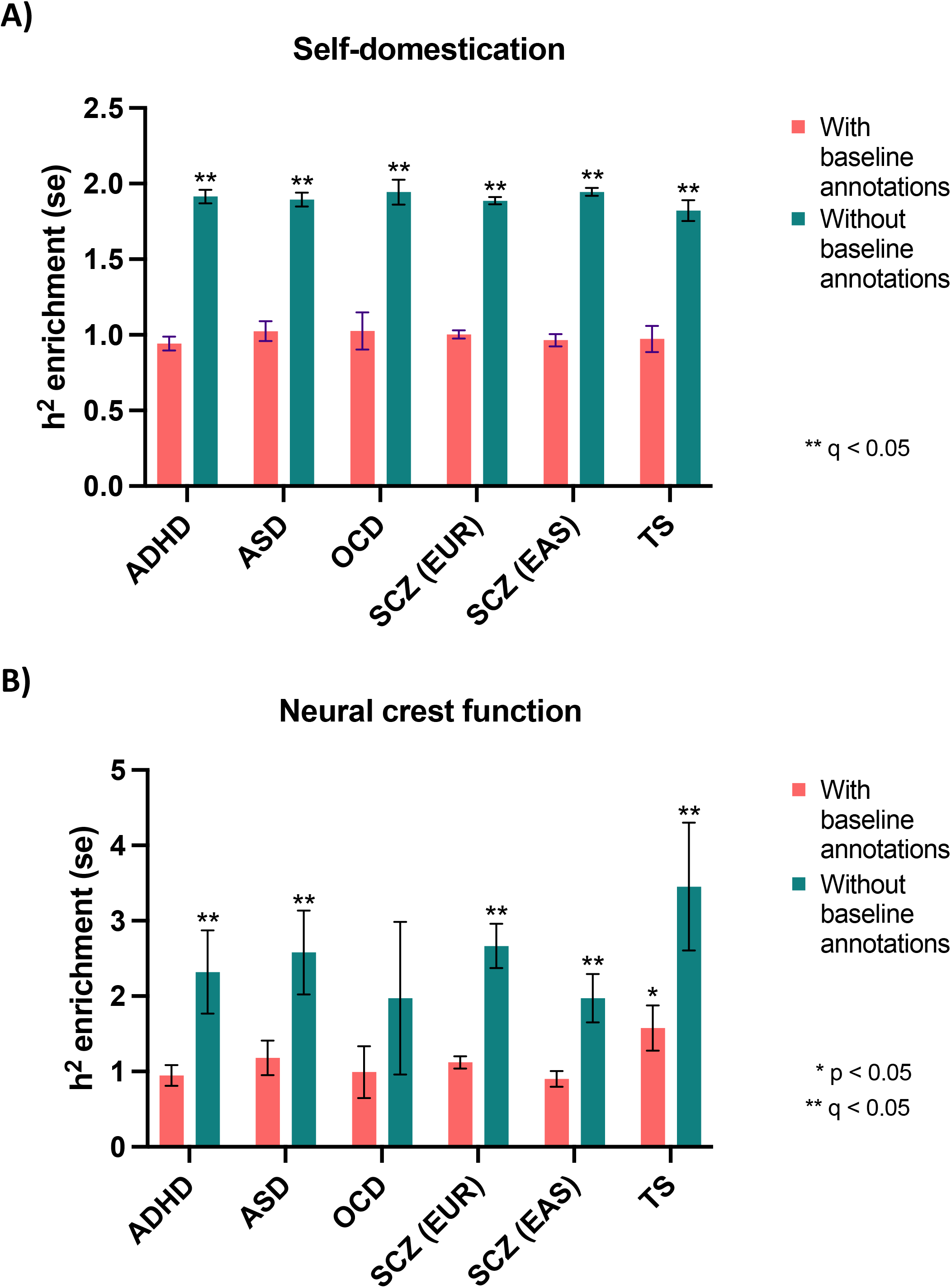
The enrichment of self-domestication (Panel A) and neural crest function (Panel B) in the SNP-based heritability (SNP-h2) of attention deficit-hyperactivity disorder (ADHD), autism spectrum disorder (ASD), obsessive compulsive disorder (OCD), schizophrenia (SCZ) and Tourette’s syndrome (TS).

However, comparing the results obtained between the two annotations, we observed different patterns. The enrichments for self-domestication annotation were similar across neurodevelopmental disorders, ranging from 0.902 (EAS SCZ, p=4.55×10^−20^) to 1.577 (TS, p=5.85×10^−5^). Conversely, a wider spectrum was present for neural crest function annotation with the highest enrichment observed for TS (enrichment=3.453, p=2.88×10^−3^) and the lowest for EAS SCZ (enrichment=1.971, p=3.81×10^−3^). The strength of the statistical significance of the enrichments was partially driven by the statistical power of the genome-wide association statistics. Indeed, since EUR SCZ is the most powerful dataset among those investigated (SNP-*h*^2^_SNP_ z=34.3, we observed the most significant results for both annotations with respect to this disorder (self-domestication: enrichment=1.122, p=2.07×10^−24^; neural crest function: enrichment=2.666, p=6.52×10^−20^).

As mentioned above, LDSC permits to model multiple annotations simultaneously. Since previous studies showed that accounting for multiple annotations can assess the independence of evolution-related enrichments with respect to other molecular properties (Pardinas, et al. 2018; Wendt, et al. 2021), we investigate whether the enrichments observed for self-domestication and neural crest function were independent of annotations related to different function and characteristics of human genome variation. Specifically, we used the baseline LDSC model that includes 95 genomic annotations (Supplemental Table 5) covering a wide range of functional categories (Finucane, et al. 2015). Considering this model that more accurately represents the complexity of the human genome, the enrichments for self-domestication and neural crest function were not significant anymore (Figure 1; Supplemental Tables 3 and 4). The only nominally significant was observed for TS with respect to neural crest function (enrichment=1.822, p=0.042). This change in the results of the annotations related to self-domestication and neural crest function was not due to a major change in the statistical power of the LDSC analysis in relation to the number of annotations tested. Indeed, no major change was observed in the enrichments for the baseline annotations with and without the inclusion of the annotations related to self-domestication and neural crest function (Supplemental Tables 5-10).

In the present study, we provide novel insights into the understanding of the potential role of self-domestication on neurodevelopmental processes. Understanding whether the cognition and behaviors of modern humans may have been shaped by processes similar to those observed in domesticated animals has been a primary focus of evolutionary biologists and paleoanthropologists. However, it is difficult to formally test this hypothesis (Wilkins 2020). A recent experimental study found genetic evidence supporting the human self-domestication hypothesis and its link with abnormal neurodevelopment through the analysis of neural-crest stem cells (Zanella, et al. 2019). Comparing paleogenetic data from modern and archaic humans, a modern-specific enrichment for regulatory changes both in *BAZ1B* (Bromodomain adjacent to zinc finger domain, 1B) and its experimentally defined downstream targets was found. Altered expression of BAZ1B has been related to dog hypersociability compared to wolves (vonHoldt, et al. 2018). The gene is deleted in Williams-Beuren Syndrome (WBS), a complex condition featuring hypersociability (Morris 2010). The study by Zanella and colleagues demonstrated that the BAZ1B, a tyrosine-protein kinase chromatin remodeler, is a master regulator of the modern human face and that the craniofacial and cognitive/behavioral traits observed in WBS due to *BAZ1B* deletion may be an entry point to the evolution of the modern human face and human prosociality (Zanella, et al. 2019). Interestingly, WBS shows increased traits of self-domestication (Niego and Benitez-Burraco 2022). As mentioned above, previous studies found preliminary evidence supporting altered human self-domestication and neural crest function in the predisposition to ASD (Benitez-Burraco, et al. 2016) and SCZ (Benitez-Burraco, et al. 2017). Nonetheless, no study tested this hypothesis systematically and with respect to other neurodevelopmental disorders. In the present study, we leveraged large-scale genome-wide information testing whether loci in domestication processes and neural crest function were enriched for the genetic predisposition to five neurodevelopmental disorders. As the mechanisms underlying self-domestication and neural crest function are complex mechanisms, our genome-wide approach is likely able to capture the involvement of multiple genes. Indeed, we found statistically significant enrichments for both annotations across the neurodevelopmental disorders investigated. However, these findings were not independent of genomic regulatory functions, characterizing molecular properties such as allele frequency distributions, conserved regions of the genome, and regulatory elements (Finucane, et al. 2015). Enhancers and conserved regions of the genome present an accelerated evolutionary rate in humans (Moon, et al. 2019). Previous studies found selective pressures on genes related to self-domestication in early humans (Theofanopoulou, et al. 2017) and Bronze Age Europeans (Benitez-Burraco, Chekalin, et al. 2021). Our study qualifies these findings. Hence, the genes we considered selected in domesticated animals seem to be under selective pressure in humans due to genomic functions independent of those that acted in the domestication processes in other animals. This is not against the hypothesis that our species went through a process of self-domestication. The genes in our analysis were indeed selected in humans and could be responsible for the emergence of domesticated features in our species. Moreover, they can contribute to cognitive and behavioral impairment due to abnormal self-domestication features in people with certain neurodevelopmental disorders, like ASD, or SCZ. Of particular interest, the enrichments for self-domestication and neural crest function annotations were the highest with respect to TS. Also, the enrichment for neural crest function remained nominally significant even when we accounted for the other 95 genomic annotations. This is particularly relevant because the control of aggression is strongly impacted in TS (Ashurova, et al. 2021). At the same time, we found that this enrichment was not independent of genomic regulatory functions. But as different genes seem to have been selected in different domesticated species, so that several gene pathways seem to lead to domestication (Hou, et al. 2020), we cannot rule out the possibility that there are some potential candidates for mammal domestication that have been selected in humans independently of genomic regulatory functions. Further studies are needed to clarify this possibility.

In conclusion, our findings do not undermine the self-domestication hypothesis of human evolution, but show that these associations are not independent of genomic regulatory features. Further studies will be needed to understand the complex interplay among self-domestication, neural crest function, genome regulatory features, epigenetic mechanisms, and brain biology in the context of human evolution and neurodevelopmental processes.

## Material and Methods

### Gene selection

The list of loci for mammal domestication encompasses 764 genes (Supplemental Table 1). This list resulted from merging genes that have been found positively selected in several domesticates, including guinea pig, pig, rat, dog, cat, cattle, domesticated fox, horse, rabbit, and sheep (Womack 2005; Trut, et al. 2009; Albert, et al. 2012; Axelsson, et al. 2013; Bellone, et al. 2013; Carneiro, et al. 2014; Montague, et al. 2014; Qanbari, et al. 2014; Schubert, et al. 2014; Wilkins, et al. 2014; Wright 2015; Cagan and Blass 2016; Marsden, et al. 2016; Zapata, et al. 2016; Benitez-Burraco, et al. 2017; Theofanopoulou, et al. 2017; Pendleton, et al. 2018). The list of NC-function loci encompasses 89 genes (Supplemental Table 2). These candidates were derived using pathogenic and functional criteria: neurocristopathy-associated genes annotated in the OMIM (Online Mendelian Inheritance in Man) database (available at http://omim.org/) (Amberger, et al. 2019), NC markers, genes that are functionally involved in NC induction and specification, genes involved in NC signaling (within NC-derived structures), and genes involved in cranial NC differentiation (Supplemental Table 11).

Using the gene locations available from MAGMA (de Leeuw, et al. 2015) (https://ctg.cncr.nl/software/magma), we selected common SNPs in reference to the human genome build 37 in two populations, Europeans (EUR) and East Asians (EAS). Gene locations were defined from transcription start site to stop site. The SNP lists were created from phase 3 of the 1000 Genomes Project (Genomes Project, et al. 2015).

### Data sources

Genome-wide association statistics informative of ADHD (N_eff_=51,306) (Demontis, et al. 2019), ASD (N_eff_=44,367 (Grove, et al. 2019), OCD (N_eff_=7,780) (International Obsessive Compulsive Disorder Foundation Genetics and Studies 2018), SCZ_EUR_ (N_eff_=126,281) (Trubetskoy, et al. 2022), SCZ_EAS_ (N_eff_=30,515) (Trubetskoy, et al. 2022) and TS (N_eff_=12,783) (Yu, et al. 2019) were obtained from previous studies of the Psychiatric Genomics Consortium (PGC). The effective sample size was calculated as recommended previously (Willer, et al. 2010). For SCZ, genome-wide association statistics were available for two ancestry groups, EUR and EAS. Unfortunately, no large-scale SCZ datasets were available for other ancestry groups and only EUR datasets were available for the other disorders investigated.

### Partitioned heritability analysis

SNP-h2 partitioning (Finucane, et al. 2015) was performed considering 95 baseline genomic annotations (Supplemental Table 5; baseline-LD model v2.2 downloaded from https://alkesgroup.broadinstitute.org/LDSCORE/) characterizing multiple molecular properties such as allele frequency distributions, conserved regions of the genome, and regulatory elements (Finucane, et al. 2015). We created additional genome-wide annotations for self-domestication and neural crest function. Binary annotations were created; SNPs in genes potentially implicated in self-domestication and neural crest function (see *Gene selection*) were assigned a “1”, and SNPs in other genes were assigned a “0”. Two sets of analyses were performed, with and without accounting for the baseline annotations to investigate if self-domestication and neural crest function enrichment are independent of the baseline annotations. The LD scores used in this analysis were generated from the 1000 Genome Project Phase 3 EUR and EAS reference panels. FDR multiple testing correction (q < 0.05) was applied accounting for the number of phenotypes and annotations tested.

## Data availability

Data used in this study are publicly available as of the date of publication.

Baseline genomic annotations: https://alkesgroup.broadinstitute.org/LDSCORE/.

LDSC heritability and partitioned heritability: https://github.com/bulik/ldsc/wiki.

Summary statistics from the PGC: https://pgc.unc.edu/for-researchers/download-results/

All original code is deposited at the repositories and is publicly available as of the date of publication. This paper does not report new code.

## Acknowledgements

The authors acknowledge support from the Horizon 2020 Marie Sklodowska-Curie Individual Fellowship from the European Commission (101028810), the National Institutes of Health (RF1 MH132337, R33 DA047527, and R21 DC018098) and One Mind, and MCIN/AEI/ 10.13039/501100011033 (grant PID2020-114516GB-I00 to ABB).

